# Comparative palatability of orally disintegrating tablets (ODTs) of Praziquantel (L-PZQ and Rac-PZQ) versus current PZQ tablet in African children: a randomized, single-blind, crossover study

**DOI:** 10.1101/605170

**Authors:** Muhidin K Mahende, Eric Huber, Elly Kourany-Lefoll, Ali Ali, Brooke Hayward, Deon Bezuidenhout, Wilhelmina Bagchus, Abdunoor M Kabanywanyi, On behalf of the Pediatric Praziquantel Consortium

## Abstract

**Background:** Praziquantel (PZQ) is currently the only recommended drug for infection and disease caused by the species of schistosome infecting humans; however, the current tablet formulation is not suitable for preschool age children mainly due to its bitterness and the size of the tablet. We assessed the palatability of two new orally disintegrating tablet (ODT) formulations of PZQ.

**Methods:** This randomized, single-blind, crossover, swill-and-spit palatability study (NCT02315352) was carried out at a single school in Tanzania in children aged 6–11 years old, irrespective of schistosomiasis infection. Children were stratified according to age group (6–8 years or 9–11 years) and gender, then randomized to receive each formulation in a pre-specified sequence. Over 2 days, the children assessed the palatability of levo-Praziquantel (L-PZQ) ODT 150 mg and Racemate Praziquantel (Rac-PZQ) ODT 150 mg disintegrated in the mouth without water on the first day, and L-PZQ and Rac-PZQ dispersed in water and the currently available PZQ 600 mg formulation (PZQ-Cesol®) crushed and dispersed in water on the second day. The palatability of each formulation was rated using a 100 mm visual analogue scale (VAS) incorporating a 5-point hedonic scale, immediately after spitting out the test product (VAS_t=0_ primary outcome) and after 2–5 minutes (VAS_t=2–5_).

**Findings:** In total, 48 children took part in the assessment. Overall, there was no reported difference in the VAS_t=0_ between the two ODT formulations (p=0.106) without water. Higher VAS_t=0_ and VAS_t=2–5_ scores were reported for L-PZQ ODT compared with Rac-PZQ ODT in older children (p=0.046 and p=0.026, respectively). The VAS_t=0_ and VAS_t=2–5_ were higher for both ODT formulations compared with the current formulation (p<0.001 for both time points). No serious adverse events were reported.

**Interpretation:** The new paediatric-friendly formulations dispersed in water were both found to be more palatable than the existing formulation of PZQ. There may be gender and age effects on the assessment of palatability.

**Funding:** This study was funded by Merck KGaA, Darmstadt, Germany and the Global Health Innovative Technology (GHIT) Fund (Grant nos. 2013–212).

**Author summary:** Schistosomiasis or Bilharzia is among top debilitating parasitic diseases in endemic developing countries. It presents in two forms of either urinary or intestinal form. The diseases’ mode of transmission is waterborne through contact with infested water. The main group being affected in developing countries are women and children due to their frequent contact with water. WHO introduced mass drug administration program whereby drugs are distributed in endemic communities to cut off the transmission of NTDs schistosomiasis included.

Praziquantel is the sole drug for treatment of all forms of Schistosomiasis currently and it has still been proven to be highly efficacious. Preventive chemotherapy program of WHO uses the same drug as a prophylactic tool to control the disease.

The biggest challenge for this drug is its availability as a 600mg tablet with a slightly bigger size and unpleasant taste, especially for younger children. This makes uneasy administering the correct dosage of drug to school children while making preschoolers totally neglected.

This study was done as swill and spit exercise (drug was not ingested) to assess the new orally disintegrating isomers of Praziquantel, L-PZQ and Rac-PZQ which have been prepared as a 150mg tablet and improved taste as compared to the existing Praziquantel formulation. Findings from 48 African children showed that both new formulations are more palatable to younger children as compared to the existing Praziquantel formulation.

These results provide evidence for further evaluation of the clinical efficacy and tolerability of the newer formulations towards the introduction of paediatric friendly Praziquantel tablets for Schistosomiasis treatment.

## Introduction

Schistosomiasis is one of the major parasitic diseases, leading to high morbidity and mortality annually in endemic countries. Globally about 200 million people are infected (mostly in the developing world), with about 700 million people worldwide at risk of being infected.^1^ Schistosomiasis is second only to malaria as the cause of severe morbidities in sub-Saharan Africa (SSA).^2^ The three main species of the Genus *Schistosoma* that infect humans are *Schistosoma haematobium*, which causes urogenital schistosomiasis, as well as *Schistosoma japonicum* and *Schistosoma mansoni*, which cause intestinal schistosomiasis.^3^ Schistosomiasis infections are usually acquired in early childhood whilst the complications tend to appear later in adulthood.^2^ The main complications are irreversible chronic liver disease, chronic kidney disease and bladder disease, which may progress to malignancies, such as splenomegaly, hepatocellular carcinoma and urinary bladder cancer.^4^

Over 123 million children are estimated to suffer from schistosomiasis globally, causing significant paediatric health problems in endemic regions, with negative impacts on child health and development.^5, 6^ This also predisposes these younger children to nutritional deficits,^7^ such as anaemia, as well as impairment in cognitive development.^5, 8^ A World Health Organization (WHO) 2016 report showed that, in 2014, out of 1.7 billion people globally who required treatment for neglected tropical diseases, including schistosomiasis, 1.1 billion were from low–middle income countries, this number accounts to 60% of their general total populations.^9^

The *Schistosoma* species *haematobium* and *mansoni* are both prevalent in Tanzania.^2^ Recent data show high prevalence in areas close to large bodies of water in sub Saharan Africa, such as Lake Victoria, bordering Kenya, Tanzania and Uganda, with prevalence close to 80%.^10–12^ Findings in 2010 showed that Tanzania had the third highest burden of schistosomiasis among sub-Saharan countries (53.3%), after Mozambique (72.1%) and Sierra Leone (56.4%).^13^ Pre-schoolers and school children have been found to be the groups most vulnerable to schistosomiasis infection.^14^

Since 2000, several programs have been set up in Tanzania to reduce schistosomiasis and other related diseases in communities with high prevalence. In 2009 the country adopted the WHO campaign program of annual and biannual mass drug administration for soil transmitted helminthic infections, including schistosomiasis, in adults and school age children.^15^

However, pre-school children are still not included in the deworming program,^16^ and WHO does not recommend the inclusion of very young children in schistosomiasis mass treatment campaigns as it could prove disruptive and unsafe due to the fact that there is no appropriate paediatric formulation of PZQ currently available.^17^

Praziquantel is the only currently available anti-schistosome drug that is recommended for infection and disease caused by the species of schistosome infecting humans. Although it is widely used for treatment, it is also highly effective when used as prophylaxis.^18^ Zwang et al (2017) showed that oral PZQ has good absorption from the gastrointestinal tract, has few reported side effects, and is well tolerated, even in preschool children.^19^ Since the 54th World Health Assembly in 2001, WHO has been using PZQ in antihelminthic deworming campaigns for mass drug administration against schistosomiasis,^20^ with high success rate in both children and adults as reported by community based surveys.^21^

Despite the wide use of PZQ, treatment compliance may be hindered by the bitter taste of the formulation and the size of the tablet.^22^ The development of a paediatric-friendly PZQ formulation that is not bitter to taste will increase the uptake of medication among young-age populations, will help to improve treatment in routine care settings, and may pave the way for preschoolers to be enrolled in deworming campaigns.

To address this public health question, the Paediatric Praziquantel Consortium was formed in July 2012 with the goal of developing an ODT formulation of PZQ for treatment of pre-school age children with schistosomiasis and obtaining regulatory approval of this formulation to allow future access. Ifakara Health Institute (IHI, Tanzania), in collaboration with the Consortium carried out this phase I clinical trial, which evaluated the palatability of two new ODT formulations of PZQ in comparison with the current available PZQ tablets (PZQ-Cesol^®^).^23, 24^

## Methods

### Study design and participants

This was a randomized, single-blind, crossover clinical trial done as a swill-and-spit palatability assessment of the new paediatric-friendly formulations of PZQ carried out in a single school on two consecutive days over one weekend.

Children were recruited from classes I–V at Umwe primary school in Rufiji district located in south-east Tanzania, where schistosomiasis is endemic. Children and their parents were notified about the trial through an open community-based recruitment process, during which investigators conducted community-based meetings, and the parents of school children and local community leaders were invited to participate during meetings on the school premises. Detailed information about the study procedures and other trial-related requirements (e.g., consent forms) were provided to those who were willing to participate, and written consent forms were also shared to Umwe primary school teachers for them to verify the information with parents and children.

Children of either gender, irrespective of having schistosomiasis or not as this was not assessed, were eligible if they were aged 6–11 years, inclusive (age was ascertained from the parents verbally and verified from a legal birth certificate or school records), they were able to comply with the protocol, and they could communicate well with the investigator in Swahili language. Children were excluded if they were not able to attend the follow-up visits, had any pre-existing condition or dietary habit that was known to interfere with their sense of smell, taste or ingestion of medication, were febrile or had a history of body temperature ≥38°, had a respiratory and/or heart rate higher than normal, had the presence of oral thrush in the 2 weeks preceding the trial, or were in any other clinical investigation for any other pharmaceutical product in the preceding 4 weeks. All parents gave their written informed consent for the child to take part and the child gave written assent that they were willing to take part and to comply with trial procedure.

Ethical clearance was obtained from the Ifakara Health Institute review board, the national Medical Research Council committee (MRCC)-NIMR, and the ethical committee of North-Western and Central Switzerland (EKNZ; approval number EKNZ2014-400). The trial protocol and the medicinal products were further approved by the Tanzanian food and drug authority (TFDA; approval number TZ15CT006). The data safety monitoring committee was established by the sponsor.

### Trial Registration

The trial was registered on ClinicalTrials.gov (NCT02315352) and in the Pan African Clinical Trials Registry (PACTR201412000959159).

### Screening

Participants were invited to come to the school with their parents, where all screening procedures, training and further recruitment were carried out. They were screened for their ability to perform the taste assessment and instructed on how to adhere to the taste trial procedures and regulations. Screening was completed in four rounds over two consecutive weeks on consecutive days over the weekend. Children were routinely assessed for their ability to hold 2 mL of juice in their mouth for 10 seconds and then spit it out as well as their ability to keep a candy in their mouth for 20 seconds without swallowing it. They were asked to properly assess and differentiate the flavors of different drinks (e.g., mango or orange juice) and to give feedback. Those who passed these preliminary tests were instructed how to use a smiley and dismal presentation of the pictorial hedonic scale to describe their feeling after every gustatory test. Children were screened until there were 48 children could be included in 4 equal-sized groups by age (6–8 years or 9–11 years) and gender, n=12 per group.

### Test procedures and medications

Test assessments were carried out within the school premises during weekends in April and May 2015. The investigation medicinal product (IMP; L-PZQ ODT, Rac-PZQ ODT or the currently available PZQ tablet-Cesol^®^ formulation) was prepared by a pharmacist in a separate room, to which the rest of the investigation team had limited access. The study pharmacist who was issuing the investigational product was the only person who was aware of the formulation given to a particular participant. On Day 1, participants assessed the palatability of the following arms in a randomized sequence: L-PZQ ODT (150 mg) disintegrated in the mouth and Rac-PZQ ODT (150 mg) disintegrated in the mouth. On Day 2, participants assessed the palatability of the following arms in a randomized sequence: L-PZQ ODT (150 mg) dispersed in water; Rac-PZQ ODT (150 mg) dispersed in water; current PZQ tablet (150 mg, 1/4 of a 600 mg tablet) crushed and dispersed in water. The test sequence was determined from a dedicated randomization list stratified by age group and gender. Participants tasted each preparation only once in each of the five periods. There was 2 hours between each assessment.

Following each swill-and-spit assessment, the participants were directed to an assessment room separate from test room. In here, clinicians administered separate questionnaires incorporating the pictorial hedonic scale for the children to report their taste opinions from each assessment period. Children were asked to place a mark along the line of the 100 mm visual analogue scale (VAS) that incorporated a 5-point hedonic scale for overall palatability. In addition, any discomfort or other occurrences in relation to the taste of the study medication (e.g., that led to the medicine being spat out) were also recorded. An open-ended questionnaire to describe the feeling in the mouth and the taste was completed for each child during the washout period between tests.

To standardize the gustatory environment and to ensure the children were not hungry between assessments, they were given breakfast 2 hours before the first assessment. Immediately after each assessment, a cracker with no smell or taste was given to all children, to remove any residual taste. Children were kept under the supervision of an investigator to ensure they did not eat anything between tests.

### Primary outcome

The first primary outcome was the difference in VAS score taken at 0 minutes immediately after spitting out (VAS_t=0_) for L-PZQ ODT disintegrated in the mouth without water versus Rac-PZQ ODT disintegrated in the mouth without water on Day 1.

### Secondary outcomes

The secondary outcome was the difference in VAS score taken at 0 minutes immediately after spitting out (VAS_t=0_) for L-PZQ ODT (150 mg) dispersed in water or Rac-PZQ ODT (150 mg) dispersed in water versus the current PZQ-Cesol® tablet (150 mg, 1/4 of a 600 mg tablet) crushed and dispersed in water on Day 2. Other secondary outcomes were the VAS scores taken at 2 to 5 minutes (VASt_2–5_) after spitting out the IMP on Day 1 and on Day 2.

### Other outcomes

Other outcomes included the recording of any discomfort or other observation was done throughout the study duration. As this was only a swill-and-spit study, no other safety profile or tolerability assessments like blood tests were evaluated. Participants were instructed not to swallow the IMP, and even in a situation of inadvertent swallowing or high buccal absorption of the 150 mg tablet, which is one quarter of a full tablet in solution, the low dose presented negligible risk of drug effect.

### Statistical analysis

A total sample size of 48 participants was considered adequate to fulfill the objective of the study. Participants were enrolled into 4 equal-sized groups of 12 according to age (6–8 years or 9–11 years) and gender. Study data were collected using paper case report forms (CRF) and then double entered into the OpenClinica^®^ open source software onsite.^25^

Data were to be analysed using absolute and relative frequencies for qualitative variables or means (standard deviation [SD]) or median (interquartile range [IQR], minima and maxima [min, max]) for quantitative variables. Descriptive analyses of outcome variables were stratified by formulation/preparation group and time of assessment. The VAS was summarized using descriptive statistics according to formulation/preparation group.

To test the palatability, it was hypothesized that Rac-PZQ ODT (without water) and L-PZQ ODT (without water) taste different (H_A1_), as opposed to the null hypothesis (H_01_) that the tastes of the two formulations are indistinguishable. This was assessed using a linear mixed model with a factor variable for the two formulations and with random intercepts at the level of the children. The second hypothesis (H_A2_) was that at least one of the two new formulations (with water) differs in taste from the current PZQ-Cesol® formulation. Here, each of the new formulations was tested against the current formulation at α=0.025. The test sequence, gender, and age of the participants were considered when testing these hypotheses.

Linear mixed modelling was also used to determine the association among VAS scores and gender and age, and to identify potential interactions of these factors with the taste of the formulations. Backward model selection was conducted guided by the Akaike information criterion (AIC). The model with the lowest AIC value that included at least the independent factors (sequence, formulation, gender, and age) was selected as the final model. P-values were considered significant at the two-sided α=0.05 level with no adjustment for multiple comparisons across models. Analysis was completed using R.^26^

## Results

### Participants

During screening, 75 children who had undergone repeated training rounds on how to adhere to study procedures were identified. Of these, 48 children were deemed eligible for participation (Figure 1), distributed in equal-sized groups according to age group and gender and randomized such that equal numbers received each test sequence in each group. The weight of the participants ranged from 14.7 to 33.1 kg, corresponding to PZQ doses of 4.5 to 10.2 mg/kg if the 150 mg dose were to be unintentionally swallowed during the process. Mean (SD) body temperature, respiratory rates and heart rates were within normal ranges before and after each assessment (Table 1).

**Figure 1.**
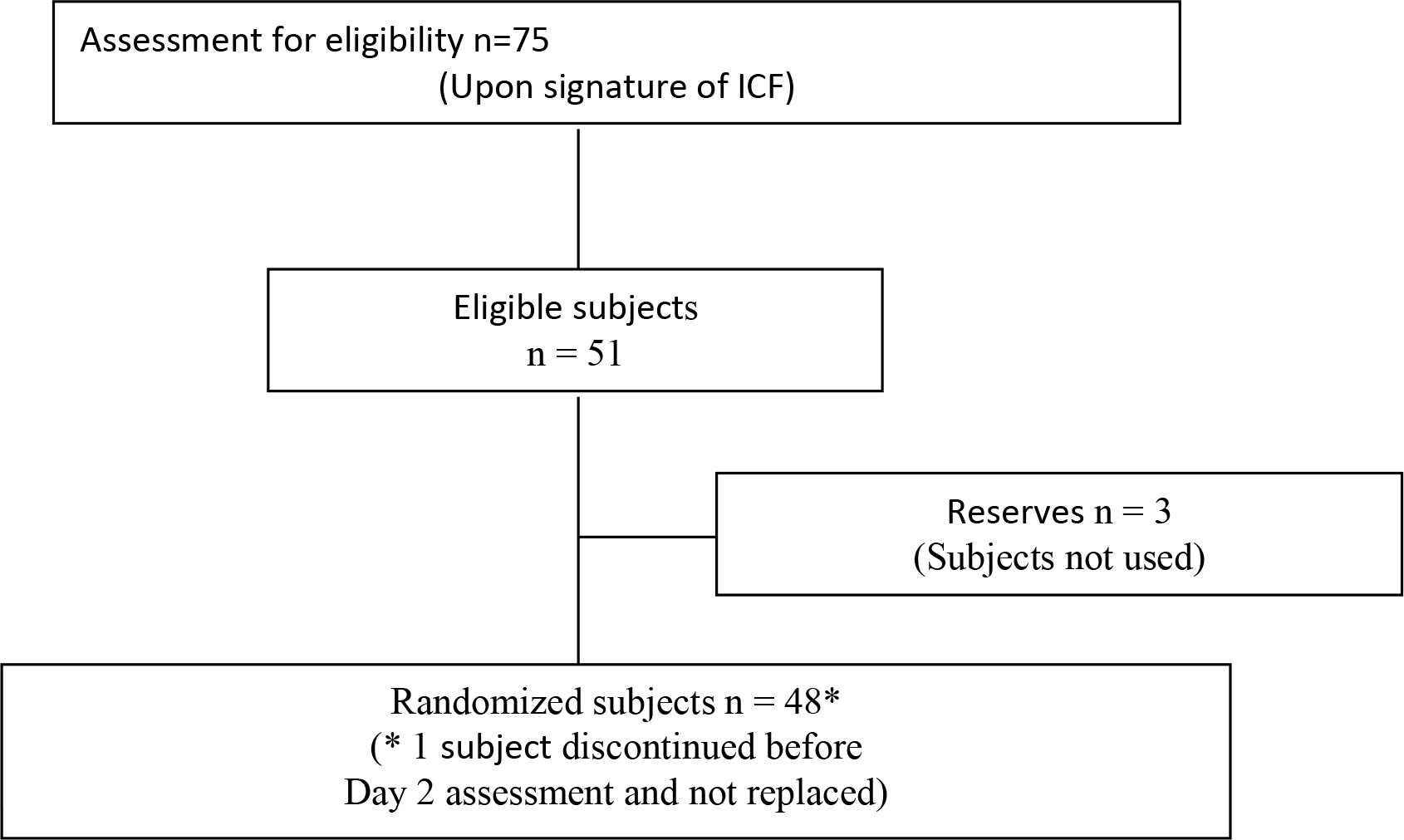
Flow diagram of participant distribution showing number of participants assessed for eligibility, enrolled, and randomized.

**Figure 2.**
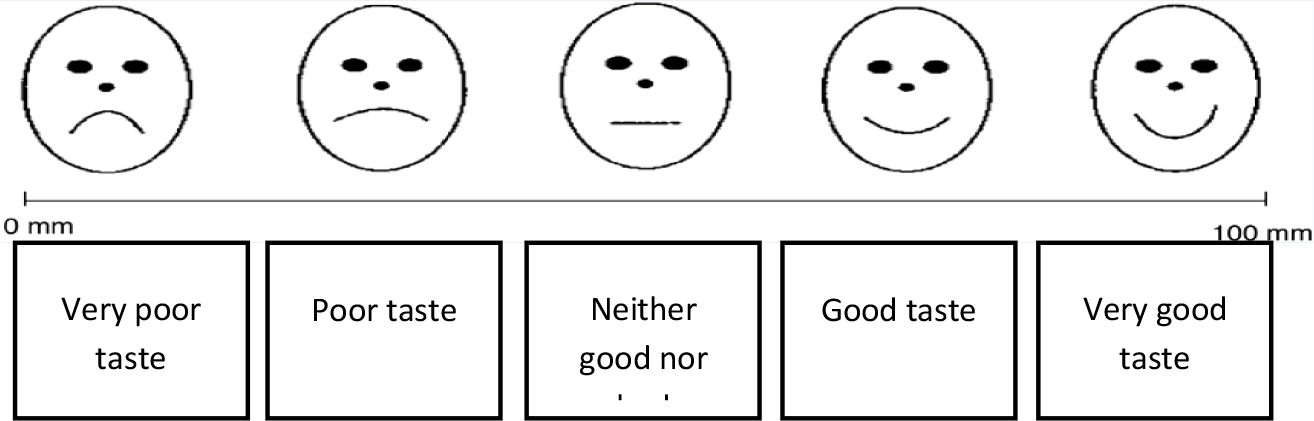
100 mm Hedonic VAS scale

**Table 1.** Vital signs by visit.

| Day of observation | N participants | Heart rate, beats per minute<br>Mean (SD) | Temperature, C°<br>Mean (SD) | Respiratory rate, breaths per minute<br>Mean (SD) |
| --- | --- | --- | --- | --- |
| At screening | 48 | 87.0 (6.23) | 36.5 (0.34) | 22.5 (4.03) |
| Day 1 before assessment | 48 | 85.1 (6.19) | 36.3 (0.24) | 19.5 (1.92) |
| Day 1 after assessment | 48 | 85.3 (5.51) | 36.5 (0.26) | 19.3 (1.33) |
| Day 2 before assessment | 47 | 85.6 (5.62) | 36.3 (0.24) | 19.5 (1.63) |
| Day 2 after assessment | 47 | 85.6 (4.93) | 36.5 (0.28) | 19.3 (1.58) |
| SD, standard deviation. |  |  |  |  |

### Primary outcome

The overall mean (SD) VAS_t=0_ scores in millimeters (both age groups and genders combined) for L-PZQ ODT and Rac-PZQ ODT without water were 49.0 (33.7) and 39.3 (28.4), respectively. The mean difference (SD) in VAS_t=0_ scores between the two formulations was not significant (9.6 [40.3]; p=0.106) (Table 2).

**Table 2.** VAS scores (mm) for L-PZQ ODT and Rac-PZQ ODT without water (Day 1) at time points 0 and 2–5 minutes.

| Formulation | At time 0 minutes |  | At time 2–5 minutes |  |
| --- | --- | --- | --- | --- |
|  | Mean (SD) | *p*-value | Mean (SD) | *p*-value |
| All participants, n=48 |  |  |  |  |
| L-PZQ ODT | 49.0 (33.65) |  | 50.7 (27.23) |  |
| Rac-PZQ ODT | 39.3 (28.43) |  | 45.1 (29.62) |  |
| Difference (L-Rac) | 9.6 (40.29) | 0.106<sup>a</sup> | 5.6 (36.15) | 0.284<sup>b</sup> |
| Among 6 - 8 year olds, n=24 |  |  |  |  |
| L-PZQ ODT | 49.6 (38.28) |  | 51.5 (25.96) |  |
| Rac-PZQ ODT | 45.8 (33.14) |  | 56.2 (28.81) |  |
| Difference (L-Rac) | 3.8 (44.11) | 0.680<sup>c</sup> | -4.7 (37.42) | 0.555<sup>c</sup> |
| Among 9 - 11 year olds, n=24 |  |  |  |  |
| L-PZQ ODT | 48.3 (29.11) |  | 50.0 (28.97) |  |
| Rac-PZQ ODT | 32.8 (21.60) |  | 34.0 (26.60) |  |
| Difference (L-Rac) | 15.5 (36.07) | 0.046<sup>c</sup> | 15.9 (32.39) | 0.026<sup>c</sup> |
| SD, standard deviation |  |  |  |  |
| <sup>a</sup>From linear mixed model adjusted for period, gender and age group. No interactions were retained in the model. |  |  |  |  |
| <sup>b</sup>From linear mixed model adjusted for period, gender, age group and interaction terms for age with gender and age with formulation. |  |  |  |  |
| <sup>c</sup>From linear mixed model adjusted for period and gender. No interactions were retained in the model. |  |  |  |  |

### Secondary outcomes

#### L-PZQ ODT and Rac-PZQ ODT without water

The overall mean (SD) VAS_t=2–5_ scores for L-PZQ ODT and Rac-PZQ ODT without water were 50.7 (27.2) and 45.1 (29.6), respectively. The overall mean (SD) difference in VAS_t=2–5_ scores between the two formulations was not significant (5.6 [36.1]; p=0.284), however there was a significant association between formulation and age group at this time point (interaction p=0.0499). The mean VAS scores for L-PZQ ODT without water did not differ from scores for Rac-PZQ ODT without water in younger children (6–8 years old). Among older children (9–11 years old), VAS scores for L-PZQ ODT without water compared with Rac-PZQ ODT without water were higher by more than 15 mm (p=0.046 and p=0.026 at time 0 and 2–5 minutes, respectively) (Table 2). The distributions of VAS_t=0_ and VAS_t=2–5_ scores by gender and age group for the ODT formulations without water are shown in Figures 3 and 4.

**Figure 3.**
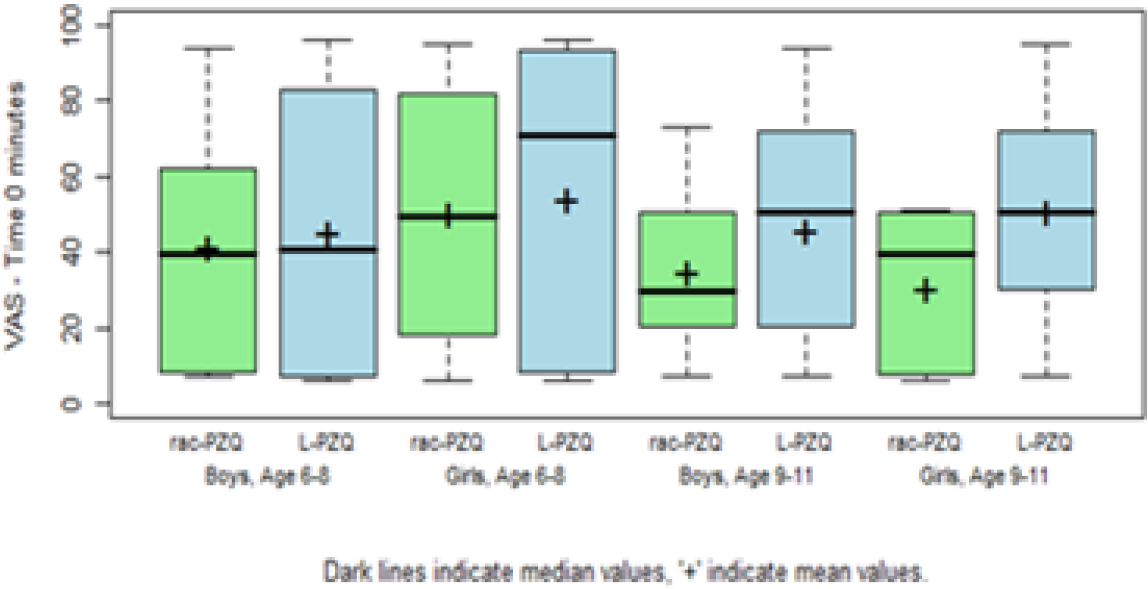
Boxplot of VAS scores (mm) for L-PZQ ODT and Rac-PZQ ODT without water at time 0 minutes, by gender and age

**Figure 4.**
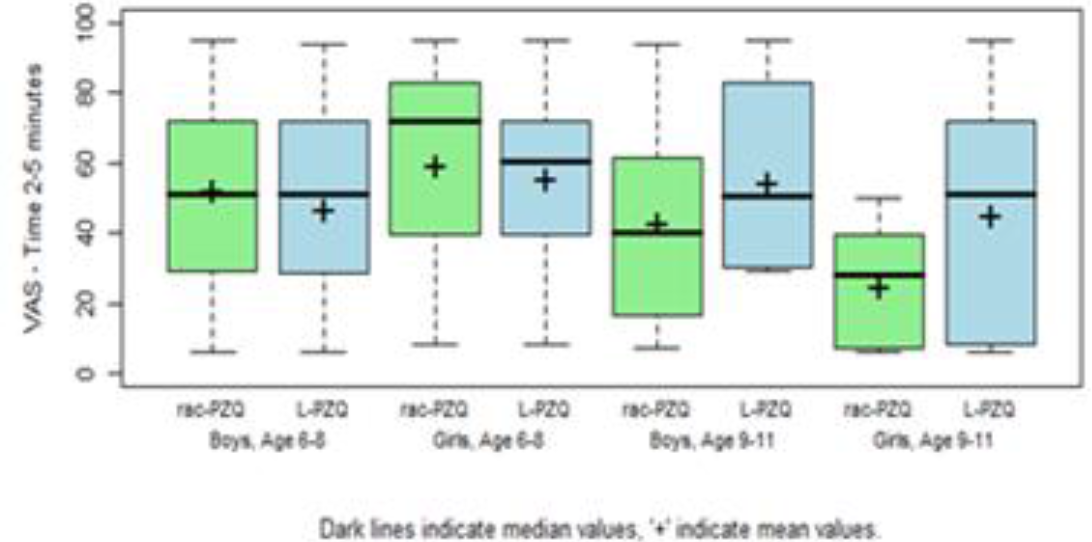
Boxplot of VAS scores (mm) for L-PZQ ODT and Rac-PZQ ODT without water at time 2-5 minutes, by gender and age

#### L-PZQ ODT, Rac-PZQ ODT and current PZQ-Cesol® with water

The overall mean (SD) VAS_t=0_ score for current PZQ-Cesol^®^ crushed in water was 20.3 (24.9) mm, while for L-PZQ ODT dispersed in water it was 67.5 (31.1) mm and for Rac-PZQ ODT dispersed in water it was 51.4 (32.4) mm. Similarly, at 2 – 5 minutes, the overall mean (SD) VAS_t=2–5_ scores were 33.5 (22.2) for current PZQ-Cesol®, 61.1 (21.2) for L-PZQ ODT and 50.2 (25.9) for Rac-PZQ ODT in water. The differences in VAS scores for L-PZQ ODT and Rac-PZQ ODT dispersed in water compared with PZQCesol^®^ were higher at both time points (p<0.001; Table 3). In addition, there was an association between the L-PZQ ODT formulation and gender (interaction p=0.004) with higher VAS_t=0_ scores for L-PZQ ODT in boys. When stratifying by gender, VAS scores for L-PZQ ODT and Rac-PZQ ODT dispersed in water were also higher than for PZQ-Cesol^®^ at each time point (p<0.01; Table 3). The distributions of VAS_t=0_ and VAS_t=2–5_ scores by gender and age group are shown in Figures 5 and 6.

**Figure 5.**
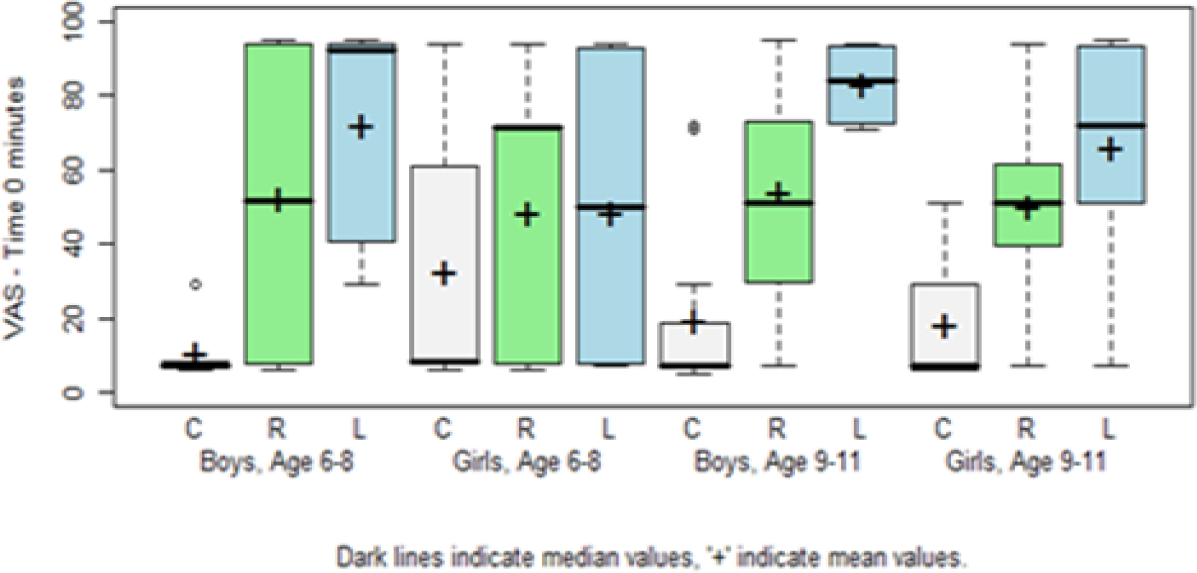
Boxplot of VAS scores (mm) for L-PZQ ODT (L) and Rac-PZQ ODT (R) dispersed with water and current PZQ-Cesol® (C) crushed in water at time 0 minutes, by gender and age.

**Figure 6.**
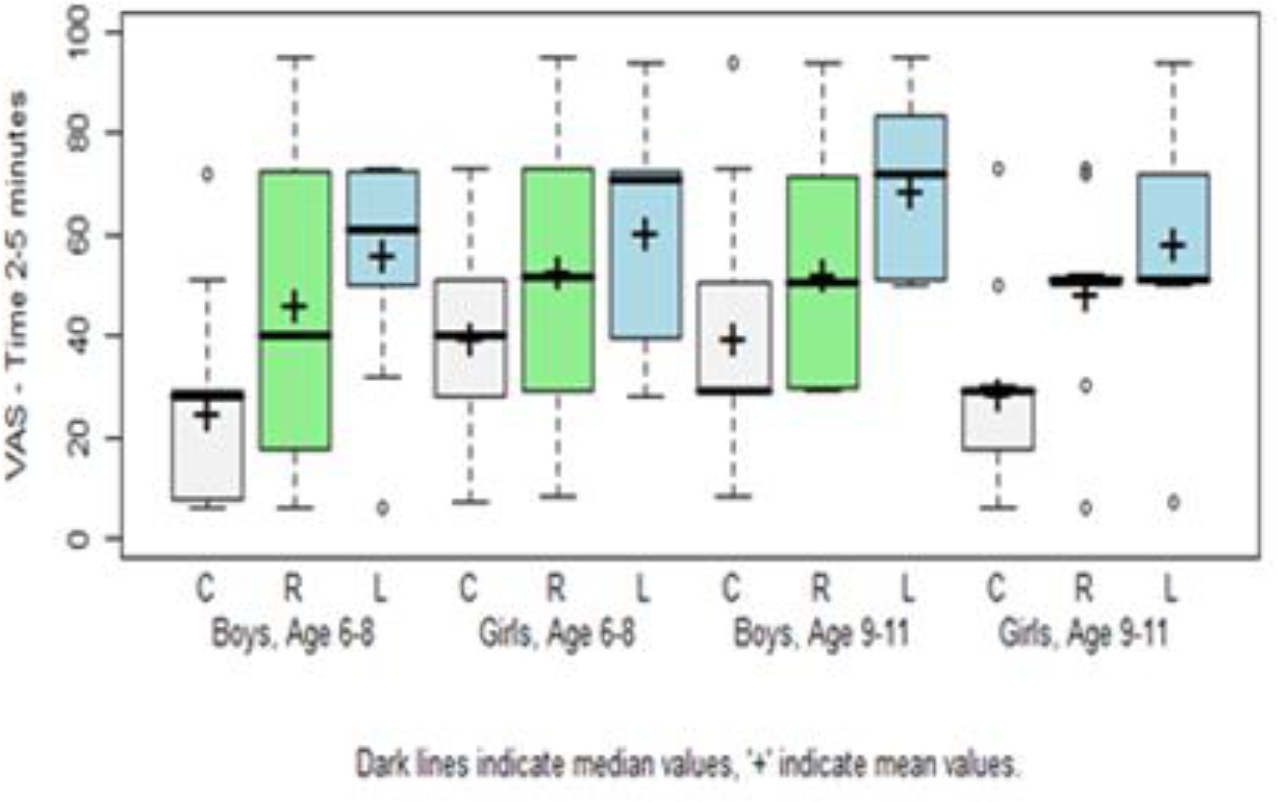
Boxplot of VAS scores (mm) for L-PZQ ODT (L) and Rac-PZQ ODT (R) dispersed with water and current PZQ-Cesol® (C) crushed in water at time 2-5 minutes, by gender and age.

**Table 3.** VAS scores (mm) from linear mixed model analysis stratified by gender for L-PZQ ODT dispersed in water and Rac-PZQ ODT dispersed in water relative to current PZQ-Cesol^®^ crushed in water, at time 0 and 2–5 minutes.

| Formulation | At time 0 minutes |  | At time 2–5 minutes |  |
| --- | --- | --- | --- | --- |
|  | Mean (SD) | *p*-value | Mean (SD) | *p*-value |
| All participants, n=47 |  |  |  |  |
| L-PZQ ODT | 67.5 (31.14) |  | 61.1 (21.19) |  |
| Rac-PZQ ODT | 51.4 (32.35) |  | 50.2 (25.92) |  |
| PZQ-Cesol® | 20.3 (24.91) |  | 33.5 (22.23) |  |
| Difference (L-Cesol®) | 47.2 (38.27) | <0.001<sup>a</sup> | 27.7 (23.58) | <0.001<sup>b</sup> |
| Difference (Rac-Cesol®) | 31.1 (38.13) | <0.001<sup>a</sup> | 16.7 (30.15) | <0.001<sup>b</sup> |
| Difference (L-Rac) | 16.1 (37.46) | 0.002<sup>a</sup> | 11.0 (27.50) | 0.008<sup>b</sup> |
| Among Boys, n=24 |  |  |  |  |
| L-PZQ ODT | 77.6 (22.02) |  | 62.5 (20.25) |  |
| Rac-PZQ ODT | 53.3 (34.23) |  | 49.5 (27.60) |  |
| PZQ-Cesol® | 15.0 (18.92) |  | 32.4 (23.64) |  |
| Difference (L-Cesol®) | 62.6 (27.23) | <0.001<sup>c</sup> | 30.1 (26.40) | <0.001<sup>c</sup> |
| Difference (Rac-Cesol®) | 38.3 (41.70) | <0.001<sup>c</sup> | 17.1 (31.43) | 0.005<sup>c</sup> |
| Difference (L-Rac) | 24.3 (31.39) | 0.001<sup>c</sup> | 13.0 (26.20) | 0.010<sup>c</sup> |
| Among Girls, n=23 |  |  |  |  |
| L-PZQ ODT | 57.0 (35.99) |  | 59.7 (22.48) |  |
| Rac-PZQ ODT | 49.4 (30.89) |  | 50.9 (24.63) |  |
| PZQ-Cesol® | 25.9 (29.34) |  | 34.6 (21.12) |  |
| Difference (L-Cesol®) | 31.2 (41.97) | <0.001<sup>c</sup> | 25.2 (20.52) | <0.001<sup>c</sup> |
| Difference (Rac-Cesol®) | 23.6 (33.26) | 0.007<sup>c</sup> | 16.3 (29.45) | 0.005<sup>c</sup> |
| Difference (L-Rac) | 7.6 (41.89) | 0.431<sup>c</sup> | 8.8 (29.23) | 0.181<sup>c</sup> |
| SD, standard deviation |  |  |  |  |
| <sup>a</sup>From linear mixed model adjusted for period, gender, age group and interaction term for gender with formulation. |  |  |  |  |
| <sup>b</sup>From linear mixed model adjusted for period, gender and age group. No interactions were retained in the model. |  |  |  |  |
| <sup>c</sup>From linear mixed model adjusted for period and age group. No interactions were retained in the model. |  |  |  |  |

#### L-PZQ ODT and Rac-PZQ ODT with water

Both the mean VAS_t=0_ and VAS_t=2–5_ for L-PZQ ODT dispersed in water were significantly higher than for Rac-PZQ ODT dispersed in water (16.1 [37.5] and 11.0 [27.5], respectively, p<0.01 for each time point; Table 3). When stratifying by gender, only VAS scores for boys remained significantly higher for L-PZQ ODT compared with Rac-PZQ ODT when dispersed in water.

### Adverse events

There were no severe adverse events observed throughout the trial period. Two (4.2%) participants reported occurrences of discomfort each during the study. One child reported abdominal discomfort on Day 2; this child had the Day 2 sequence of Rac-PZQ ODT/current PZQ-Cesol^®^/L-PZQ ODT. One child reported headache on Day 2; this child had the Day 2 sequence of Rac-PZQ ODT/L-PZQ ODT/PZQCesol^®^. One child (2.1%) presented with malaria, which was constitutionally considered an adverse event before tasting on Day 2. This child was removed from the study and treated with an appropriate antimalarial medication; she was followed up and was not replaced in the study with another child.

## Discussion

When the two ODT formulations were tasted without water, mean VAS scores were acceptable in the “Neither good nor bad” range of the hedonic VAS scale and the VAS score for L-PZQ ODT did not differ from that for Rac-PZQ ODT. Overall, the between-participant variability was considerable, especially for age and gender. Separate analyses for the two age groups revealed that L-PZQ ODT without water was significantly more palatable than Rac-PZQ ODT without water in children aged between 9 and 11 years. These results therefore provide some evidence for the better palatability of L-PZQ ODT without water over Rac-PZQ ODT without water in older children. This was not observed in younger children. The age differences remain an important factor to accurately report on the palatability of each preparation without water.

When tasted with water, both ODT formulations were clearly more palatable than the current PZQ-Cesol® crushed in water. Similar to without water, the mean VAS scores of the ODT formulations were in the “Neither good nor bad” range while the PZQ-Cesol® mean VAS scores were in the “Poor Taste” part of the hedonic VAS scale. Palatability reported immediately after spitting out the IMP by boys differed from what was reported by girls; boys reported higher for L-PZQ ODT and lower for PZQ-Cesol®. However, separate analyses by gender still revealed significantly higher scores for both the new formulations dispersed in water compared with the current PZQ-Cesol® formulation. It is of note that LPZQ ODT dispersed in water was more palatable compared with Rac-PZQ ODT dispersed in water at both time points for boys but not for girls. Thus, gender continues to be an important factor along with age to consider when reporting palatability. These findings may be important for the future development of the drug, which could be given as a dispersible tablet in the mouth or dispersed in water.

Our results support further development of the ODT formulations. This study was limited to children aged 6 years and older; pre-school children were not included as they are often shy, reluctant to participate and unable to communicate their feelings and preferences.^27^ However, it is reasonable to assume that the results can be extrapolated to the younger age group, who are commonly more sensitive to bitter tasting drugs compared with older children and adults.^28^ Although some parents were not familiar with the drug and the trial process before the trial started, the trial execution was a success. When the PZQ paediatric friendly formulation becomes available, a solid access pathway needs to be in place, to allow an efficient delivery ^5^ and monitoring of the drug into current healthcare setups to meet the need for African children to have better access to medication. Effective early treatment is possible, thereby preventing the substantial immune-mediated effects of schistosomiasis infection.^3, 24^

## Conclusion

In conclusion, this trial has shown that the overall palatability of both L-PZQ ODT and Rac-PZQ ODT was higher than that of PZQ-Cesol^®^ when the drugs were dispersed in water in this study population of African children. The superiority of L-PZQ ODT over Rac-PZQ ODT without water could not be confirmed; however, L-PZQ ODT dispersed in water was more palatable than Rac-PZQ ODT dispersed in water. Some evidence supporting an age effect and gender effect on palatability was also observed.

## Contributors

MKM and AMK supervised the clinical study and drafted the manuscript. AA, BH and AMK analysed the data. All authors contributed to the data interpretation and drafting of the manuscript. All authors read and approved the final manuscript.

## Acknowledgments

On behalf of the Paediatric Praziquantel Consortium we acknowledge the support given from various parties in making this work possible, the funders Merck KGaA (Darmstadt, Germany) and GHIT Fund for financial assistance, Rufiji district council for allowing investigators to conduct the study and lastly parents, children and teachers of Umwe primary school for their participation.

## Supporting Information

S1 file: Taste study Protocol

S2 file: Consort checklist taste study

